# A common Alu element insertion in the 3’UTR of *TMEM106B* is associated with an increased risk of dementia

**DOI:** 10.1101/2023.09.07.555370

**Authors:** Alana Rodney, Kul Karanjeet, Kellie Benzow, Michael D. Koob

## Abstract

Sequence variants in *TMEM106B* have been associated with an increased risk of developing several different types of dementia. As part of our efforts to generate a set of mouse lines in which we replaced the mouse *Tmem106b* gene with a human *TMEM106B* gene comprised of either a risk or protective haplotype, we conducted an in-depth sequence analysis of these alleles. We identified an AluYb8 insertion in the 3’ untranslated region (3’UTR) of the *TMEM106B* risk haplotype. We analyzed transcribed *TMEM106B* sequences using RNA-seq data available through the AD Knowledge portal and full genome sequences from the 1000Genomes databases and found that every risk allele analyzed shares this same AluYb8 insertion. The allele frequency of the risk haplotype ranges from 26% to 60% in the populations examined, and the balance of the alleles in each population is the protective haplotype. The primary sequence features of both haplotypes are distinct from the variants found in other primates. We conclude that the risk haplotype arose early in human development with a single Alu-insertion event within a unique haplotype context. Together these two non-ancestral allele variant types now appear to constitute the vast majority of the *TMEM106B* alleles but neither has come to fully predominate in any modern human population.

## INTRODUCTION

A genome-wide association study published in 2010 identified a haplotype block containing the gene encoding lysosomal type II transmembrane protein 106B (*TMEM106B*) as a susceptibility locus for Frontotemporal lobar degeneration (FTLD) with TAR DNA-binding protein (TDP-43) inclusions (FTLD-TDP) ^1^. The top tagging SNP found in this study was rs1990622, with the minor G allele reducing the risk of developing FTLD-TDP dementia and the major A allele increasing risk. Association studies published the following year confirmed that the *TMEM106B* locus modified the risk of developing FTLD-TDP in patients with mutations in the gene encoding granulin (GRN), with the risk allele decreasing the age at onset by an average of 13 years, and identified a coding SNP (rs3173615) in perfect linkage disequilibrium with rs1990622 that changes a single amino acid in the minor protective form of TMEM106B protein (T185S)^2^. A study published in 2014^3^ reported that *TMEM106B* haplotypes also modify the risk of developing dementia in patients with the hexanucleotide repeat expansion mutation in C9ORF72 but did not modify the risk of developing motor neuron disease. More recent studies have found that TMEM106B haplotypes modify the rate of cognitive decline in Parkinson Disease^4^ and impact the severity of neuropathology found in postmortem brain tissue from patients with CTE ^5^ and Limbic-predominant age-related TDP-43 encephalopathy with hippocampal sclerosis pathology (LATE-NC + HS) ^6^.

The functional variant(s) in the *TMEM106B* haplotypes have not yet been conclusively identified. To date, rs3173615 is the only coding variant identified in *TMEM106B*. A study found that the risk T185 TMEM106B protein isoform is degraded more slowly than the protective S185 isoform, leading to a two-fold increase of T185 TMEM106B protein levels without impacting mRNA expression^7^. An alternative potential functional variant identified in part on *TMEM106B* expression studies in lymphoblastoid cell lines predicts that the differences in the non-coding SNP rs1990620 (very near rs1990622) differentially impacts chromatin structure and the transcription of this gene ^8^.

Several recent studies evaluating brain tissue for fibrous TDP-43 instead found fibers made entirely or partially of TMEM106B^9-11^. TMEM106B has been shown to regulate lysosomal function^12^, suggesting a role for lysosomal pathways in the differential impacts of these two haplotypes on dementia risk and in its direct association with TDP-43 protein accumulations. Surprisingly, recent work has also shown that the TMEM106B protein can be present on the cell surface and can mediate SARS-CoV2 infection in cells^13^.

Additional experimental models are needed to support the types of detailed mechanistic studies that will help to conclusively identify the functional variant(s) in these haplotypes and to elucidate at the molecular level precisely how TMEM106B impacts neuropathology. We are working to develop a set of mouse models in which we replaced the mouse *Tmem106b* gene with a human *TMEM106B* gene comprised of either a risk or a protective haplotype. As part of these efforts, we fully sequenced the two TMEM106B genomic alleles to be introduced into these mouse lines and analyzed the sequence differences between these risk and protective haplotypes.

## RESULTS

### TMEM106B Risk and Protective Haplotype Sequencing and Analysis

We used PCR and limited sequencing of polymorphic regions to identify the BAC clones RP11-960B15 and CH17-22M18 as containing *TMEM106B* risk and protective haplotypes, respectively, and then used NGS to obtain the full sequence of the *TMEM106B* region in these two clones. We analyzed and compared the syntenic 97kb region from each sequence (Chr7:12185253 −12282003, assembly GRCH38.p14) that we have incorporated into our *TMEM106B*-Gene Replacement (*TMEM106B*-GR) mouse lines. The SNP and INDEL differences between these two haplotype sequences and the reference genome are listed in **Supplementary File S1** and are summarized for the transcribed region of *TMEM106B* in **Figure 1a**. We found that the BAC RP11-960B15 risk haplotype was essentially identical to the reference genomic sequence 38.p14, which is also a risk haplotype, with only two single nucleotide insertions identified (**Fig. 1a**). On the other hand, we found that the CH17-22M18 protective haplotype was significantly different than the reference risk haplotype, with 193 SNP and 23 INDEL variants identified. By far the most striking sequence difference that we identified between the risk and protective haplotypes, however, is the insertion of a 316bp AluYb8 element in the risk haplotypes that is not present in the protective haplotype analyzed (**Fig. 1a** and **Fib. 1b**). As shown in **Fig. 1b**, this Alu element is inserted into the 3’ UTR of the longest *TMEM106B* isoform, and, as is typical of this type of insertion event, is flanked by a short, direct repeat of the insertion site sequence.

**Figure 1.**
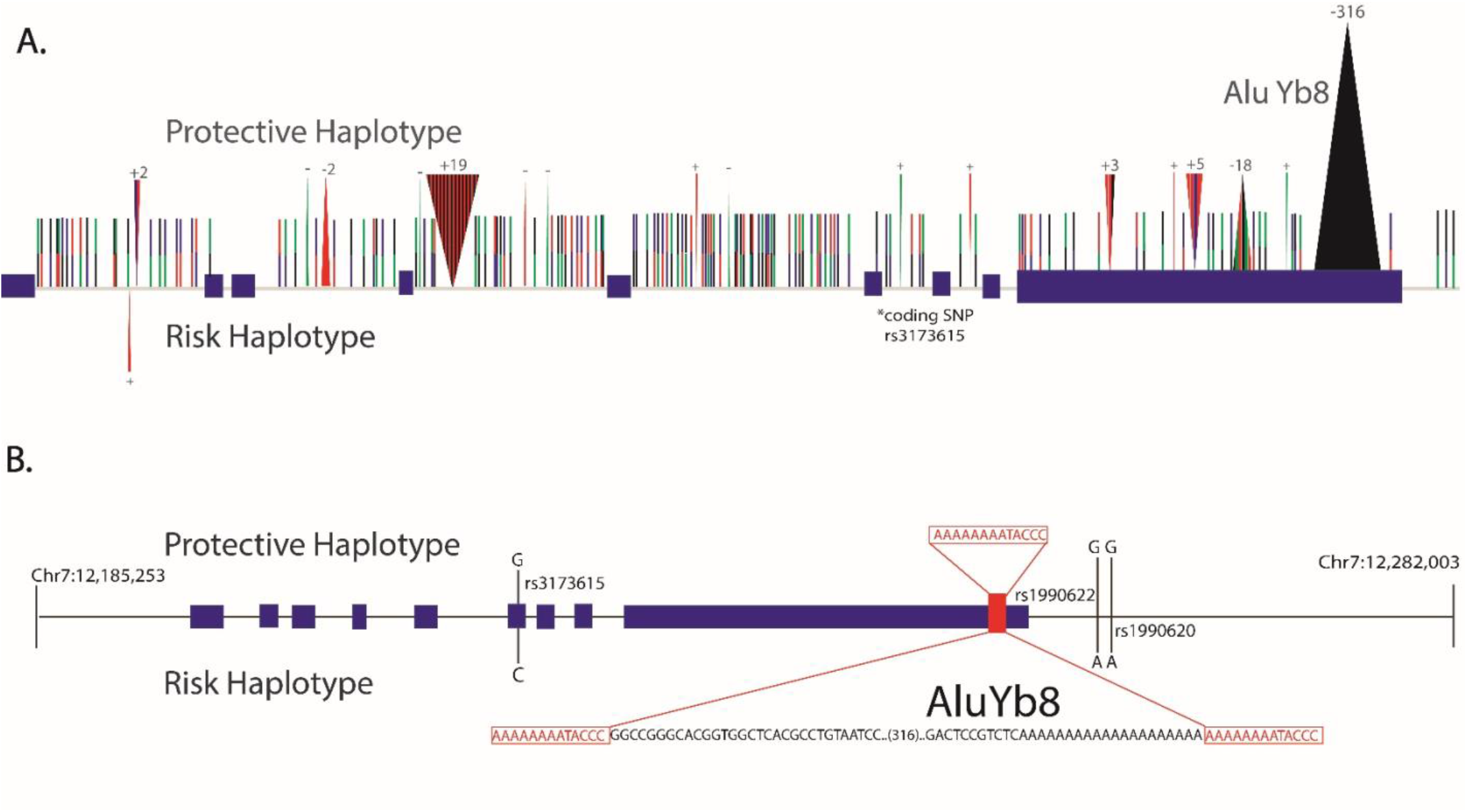
*TMEM106B* risk and protective haplotype characterization. 1A) Visual representation of variants identified in the risk and protective haplotypes within the transcribed region of TMEM106B as compared to the genome reference sequence (GRCH38.p14). Exons in the TMEM106B gene are indicated by purple boxes. 1B) Visual representation of the entire region of the risk and protective haplotypes analyzed (Chr7:12185253 −12282003,. GRCH38.p14). A 13bp direct repeat duplication at the AluYb8 insertion site is shown in red, with the head and tail ends of the Alu element in black letters. Significant SNP polymorphisms between the two haplotypes are shown. Red=A, Black=G, Green=T, Blue=C. Wedges=INDELs (# of bases added or lost is shown), Lines=SNPs

### Comparison to non-human primate haplotypes

We used BLAST analysis to non-human primate genome sequences (e.g., chimpanzee, pygmy chimpanzee, and western gorilla) and found that none shared the *TMEM106B* AluYb8 insertion. We found that the risk haplotype shared the conserved ancestral sequences at tagging SNP rs1990622-A and coding SNP rs3173615-C, but that the protective haplotype shares the putative functional SNP rs1990620-G sequence with non-human primates.

### Human RNA-seq Analysis

We digitally genotyped samples from a Mayo RNA-seq study available the AD knowledge portal based on tagging SNP rs1990622 and coding SNP rs3173615. We identified fifteen *TMEM106B* data sets homozygous for the protective haplotypes and 15 sets homozygous for the risk haplotype samples and evaluated these sequence sets for the Alu insertion sequence in *TMEM106B* (**Table 1)**. We found that all the risk haplotype data sets contained the AluYb8 insertion identified in our BAC clone haplotype, and that all the reads from the 15 protective haplotype samples contained the insertion site sequence without the Alu insertion.

**Table 1.** Human Mayo RNAseq Data. RNAseq data from 30 patients with various neurological disorders were genotyped based on the coding SNP rs3173615-G and tagging SNP rs1990622-G. All risk haplotype samples were homozygous for the AluYb8 insertion. AD=Alzheimer’s, CON=control, PSA= Progressive Supranuclear Palsey, PA= Parkinson’s.

| Table 1. Human MayoRNA Seq |  |  |  |
| --- | --- | --- | --- |
| Sample Number | Disease Phenotype | rs3173615 | ALU |
| 1015 | AD | CC | +/+ |
| 11265 | CON | CC | +/+ |
| 11289 | CON | CC | +/+ |
| 11300 | PSA | CC | +/+ |
| 11311 | CON | CC | +/+ |
| 11357 | PSP | CC | +/+ |
| 11371 | UNK | CC | +/+ |
| 11394 | PSA | CC | +/+ |
| 11424 | PA | CC | +/+ |
| 11440 | PSA | CC | +/+ |
| 11449 | PSA | CC | +/+ |
| 11249 | PSA | CC | +/+ |
| 11321 | CON | CC | +/+ |
| 11398 | PSA | CC | +/+ |
| 11497 | AD | CC | +/+ |
| 11469 | CON | GG | -/- |
| 1203 | AD | GG | -/- |
| 1302 | CON | GG | -/- |
| 1052 | AD | GG | -/- |
| 11322 | CON | GG | -/- |
| 11372 | PSA | GG | -/- |
| 11396 | CON | GG | -/- |
| 11410 | UNK | GG | -/- |
| 1929 | UNK | GG | -/- |
| 732 | AD | GG | -/- |
| 894 | AD | GG | -/- |
| 966 | AD | GG | -/- |
| 1104 | AD | GG | -/- |
| 11457 | PSA | GG | -/- |
| 1275 | AD | GG | -/- |

### Global Population Analysis

We randomly selected a set of whole genome sequencing data from the 1000 Genomes database from individuals from a range of geographically diverse populations. We found that four genomes were homozygous for the risk haplotype, five genomes that were homozygous for the protective haplotype, and 12 genomes that were heterozygous for the *TMEM106B* risk allele (**Table 2)**. We found that none of the five genomes homozygous for the protective haplotype contained the Alu insertion, that all four the genomes homozygous for the risk allele only contained sequence reads with the Alu insertion, and as expected, found the Alu insertion in half of all sequence reads from all 12 heterozygous samples.

**Table 2.**
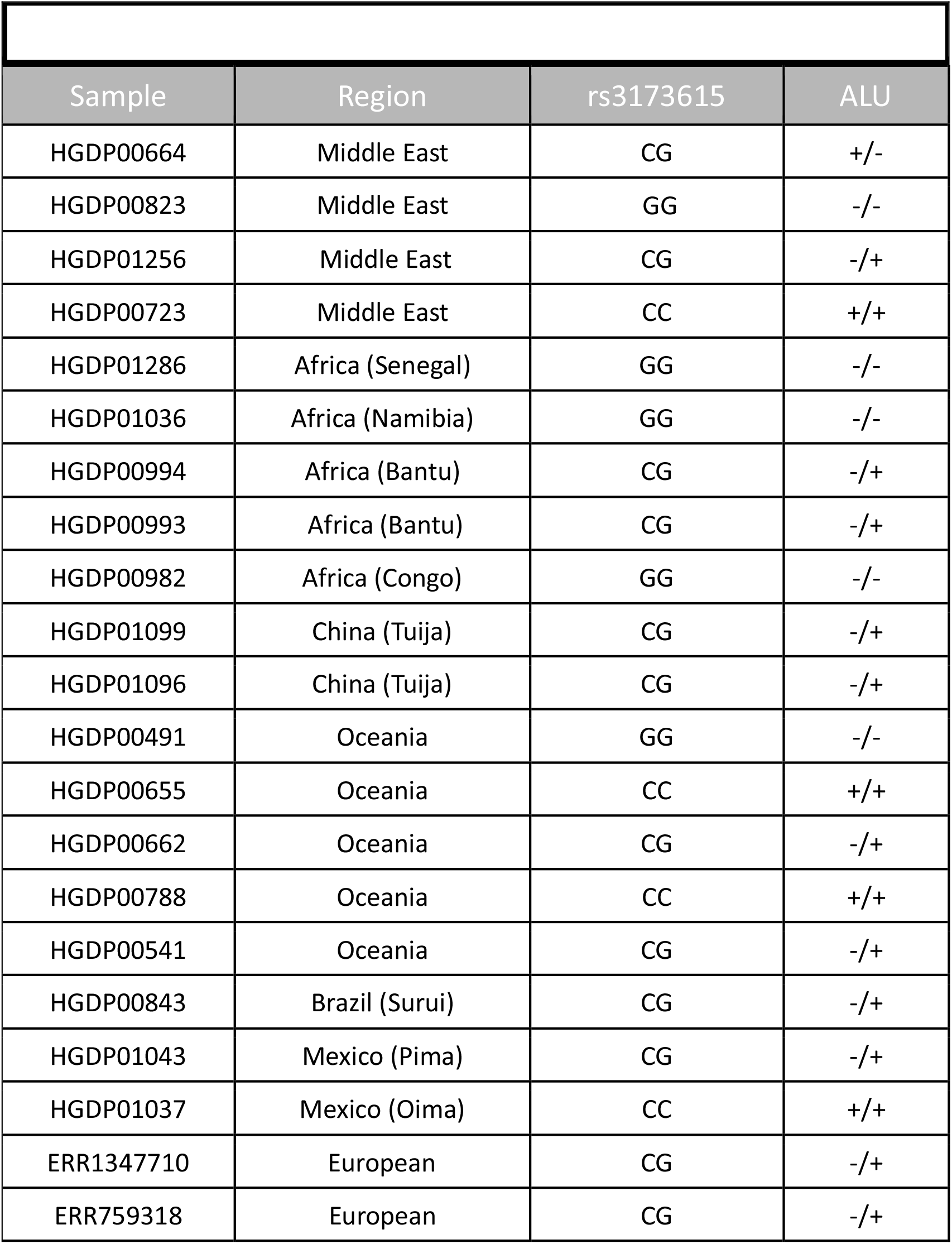
1000 Genome Dataset. Twenty-one genomes in the 1000 genome dataset from diverse populations were analyzed. Samples were genotyped using SNP rs3173615. The AluYb8 insertion was found in all risk TMEM106B haplotypes, and not in protective haplotypes.

| Sample | Region | rs3173615 | ALU |
| --- | --- | --- | --- |
| HGDP00664 | Middle East | CG | +/- |
| HGDP00823 | Middle East | GG | -/- |
| HGDP01256 | Middle East | CG | -/+ |
| HGDP00723 | Middle East | CC | +/+ |
| HGDP01286 | Africa (Senegal) | GG | -/- |
| HGDP01036 | Africa (Namibia) | GG | -/- |
| HGDP00994 | Africa (Bantu) | CG | -/+ |
| HGDP00993 | Africa (Bantu) | CG | -/+ |
| HGDP00982 | Africa (Congo) | GG | -/- |
| HGDP01099 | China (Tuija) | CG | -/+ |
| HGDP01096 | China (Tuija) | CG | -/+ |
| HGDP00491 | Oceania | GG | -/- |
| HGDP00655 | Oceania | CC | +/+ |
| HGDP00662 | Oceania | CG | -/+ |
| HGDP00788 | Oceania | CC | +/+ |
| HGDP00541 | Oceania | CG | -/+ |
| HGDP00843 | Brazil (Surui) | CG | -/+ |
| HGDP01043 | Mexico (Pima) | CG | -/+ |
| HGDP01037 | Mexico (Oima) | CC | +/+ |
| ERR1347710 | European | CG | -/+ |
| ERR759318 | European | CG | -/+ |

## DISCUSSION

In this study we present our findings from an in-depth sequence characterization of the pair of *TMEM106B* protective/risk haplotypes that we have introduced into our matched set of *TMEM106B*-GR mouse lines. We sequenced and analyzed these protective and risk haplotypes from BAC clones and identified an AluYb8 insertion in the 3’ UTR of *TMEM106B*. We did not find this Alu insertion sequence in the primate sequences that we analyzed. This finding together with the fact that the other major *TMEM106B* haplotype does not contain the Alu sequence strongly suggests that this insertion event occurred after the divergence from our last common primate ancestor. We evaluated transcribed *TMEM106B* sequences and full genome sequences from a range of global populations and consistently saw the Alu insertion in the risk haplotype sequence and absent from the protective. Allele frequency of the risk SNPs rs199062-A and rs3173615-C range from 26% to 60% in global populations (**Table 3)**. We note that the allele frequencies for these two SNPs are essentially identical in all populations except those identified in this study as “African”, suggesting that recombination may have occurred between these two SNPs in some of the haplotypes in that cohort. We conclude that the Alu insertion in the risk haplotype occurred early in human development in a single molecular event and is now one of the two major TMEM106B haplotypes in all populations globally. Each of these two *TMEM106B* haplotype alleles is likely to offer a significant selective evolutionary advantage over other ancestral variants, but neither has come to fully predominate in any modern human population.

**Table 3.**
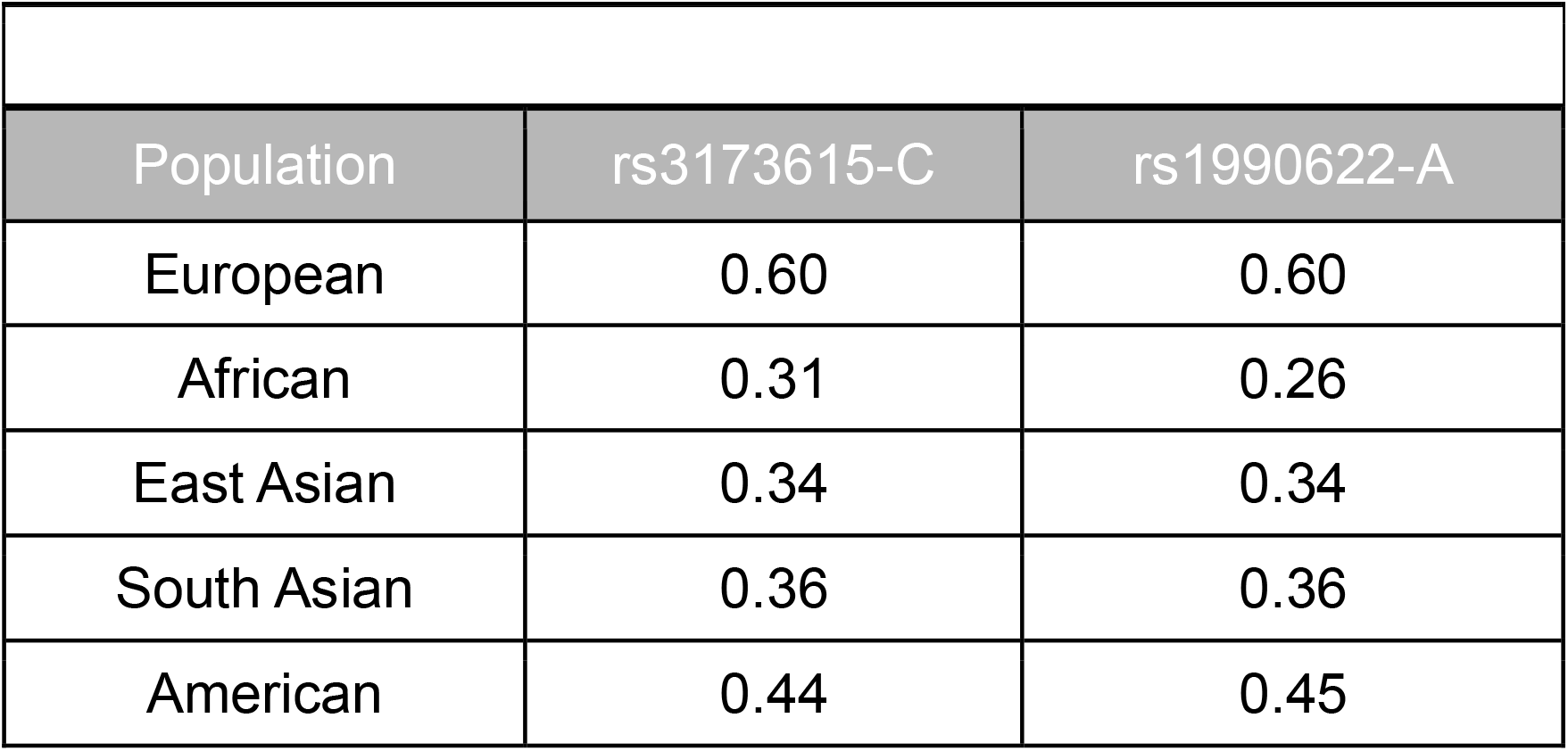
Global Allele Frequency. Allele frequency in global populations of risk alleles rs3173615-C and rs1990622-A identified in the 10000Genomes project data^21^.

| Table 3. Global Allele Frequency |  |  |
| --- | --- | --- |
| Population | rs3173615-C | rs1990622-A |
| European | 0.60 | 0.60 |
| African | 0.31 | 0.26 |
| East Asian | 0.34 | 0.34 |
| South Asian | 0.36 | 0.36 |
| American | 0.44 | 0.45 |

Although we do not yet have evidence that the AluYb8 insertion in the 3’ UTR of the risk haplotype has a functional impact on the *TMEM106B* gene in this haplotype, the 3’UTR region of genes often plays a role in transcriptional regulation^14^ and this insertion may impact the expression of this gene. Alu elements are one of the most common transposable elements in the human genome. The AluY subfamily is most recently integrated in the human genome and makes up less than 10% of the Alu insertions in the human genome^15^. Alu insertions often cause disease by disrupting a coding region or a splice signal. For example, Alu insertions in *BRCA1* can cause hereditary breast cancer^16^, and Alu insertions in *GLA* can cause cardiac diseases^17^.

The *TMEM106B*-GR mouse model set we are developing will allow us to directly evaluate the molecular impacts of the AluYb8 insertion that we identified and report here, as well as those of the other putative functional sequence variations. One of the two *TMEM106B*-GR lines that we have made to date carries the risk haplotype in place of the syntenic *Tmem106b* region of the mouse genome, and the other matched lines carries the protective haplotype (**Fig. 1**). We are currently generating additional matched lines in which we modify the sequence in the protective haplotype by a) inserting the AluYb8 sequence, or b) converting the rs3173615 SNP from G to C, and will assess the impacts of these sequence variants on TMEM106B endophenotypes in these lines.

## METHODS

### BAC Sequencing, Variant Calling and Visualization

BAC clones RP11-960B15 and CH17-22M18 containing the *TMEM106B* risk and protective haplotypes, respectively, were sequenced by the University of Minnesota Genomics Center using Illumina NovaSeq6000 with 2 x 150 bp reads. Raw sequences were mapped to the GRCh38/h38 human genome reference using Burrows-Wheeler Aligner (BWA) v0.7.17^18^, and PCR or optical duplicates were parked using Picard tools v2.19.9 (http://broadinstitute.github.io/picard/). These files were further processed with GATK v4.0.1^19^ for base quality score recalibration and variant calling **FS1**. Variants were filters using standard filtering parameters. For SNPs, the filtering criteria were QD < 2.0, FS > 60.0, SOR > 3.0, ReadPosRankSum < − 8.0, MQ < 40.0, and MQRankSum < − 12.5. For indels, the filtering criteria were QD < 2.0, FS > 200.0, SOR > 10.0, and ReadPosRankSum < − 20.0. Integrative genome viewer was used visually analyze bam files, variant calling files, and the ALU insertion in the 3’ untranslated region of the risk haplotype.

### Human RNAseq Analysis

We analyzed whole transcriptome data from the Mayo Clinic Alzheimer’s Disease Genetics Studies from the AD Knowledge portal to determine if the AluYb8 insertion was also present in diseased brains. This study contains RNA-sequencing data from cerebellum and temporal cortex samples from North American Caucasian subjects with neuropathological diagnosis of AD, progressive supranuclear palsy, pathologic aging, or elderly controls without neurodegenerative diseases^20^. Only temporal cortex samples that were homozygous for the risk or protective at SNP rs3173615 and rs1990622 were included in our analysis. Using Integrative Genome Viewer, we visualized each RNAseq bam file. BAM files were analyzed for protective or risk coding SNP and the presence or absence of the Alu insertion in the 3’ UTR region.

### Global Population Analysis

The 1000 Genomes project data set^21^ was accessed on March 15, 2023. High coverage WGS samples were the only samples included in our analysis. We genotyped these samples based on rs3173615 using IGV. Our sample set included four genomes homozygous risk, five genomes homozygous for the protective haplotype and 12 genomes that were heterozygous for these haplotypes.

## ACKNOWLEDGEMENT

The work reported here was supported by the NIA and the NINDS of the National Institutes of Health under award numbers R61/R33NS115089 and RF1AG079125.

